# Strong whole life-cycle inbreeding depression in *Daphnia magna* enhanced by partial asexuality

**DOI:** 10.1101/2020.10.16.343095

**Authors:** Valentina G. Tambovtseva, Anton A. Zharov, Christoph R. Haag, Yan R. Galimov

## Abstract

Inbreeding depression is a key factor in the evolution of mating strategies and breeding systems across the eukaryotic tree of life. Yet its potential impact in partially asexual species has only received little attention. We studied inbreeding depression in the cyclical parthenogen *Daphnia magna* by following mixtures of inbred and outbred genotypes from an early embryonic stage through hatching to adulthood and then across several asexual generations. We found that, across asexual generations, the frequency of inbred genotypes strongly and constantly decreased, until the experimental populations were almost entirely made up of outbred genotypes. The resulting estimate of inbreeding depression across the entire life cycle was almost 100 %, much higher than previous estimates for this and other similar species. Our results illustrate that the magnitude of inbreeding depression may be severely underestimated in studies that use fitness components or proxies instead of compound fitness estimates across the entire life, as well as in experimental studies with substantial pre-experimental mortality. More generally, our results suggest that inbreeding depression may play an important role in the evolution of partially asexual life cycles because clonal reproduction maintains inbreeding levels, and hence the negative effects of inbreeding accumulate across subsequent asexual generations.

## INTRODUCION

Inbreeding depression, the reduced fitness of offspring resulting from mating between related parents, is a major concern for the conservation of endangered species and a key factor shaping the evolution of life-history strategies and breeding systems (Charlesworth and Charlesworth 1987; Keller and Waller 2002; Hedrick and Garcia-Dorado 2016). Inbreeding depression is usually assessed by correlating some measure of fitness with the inbreeding coefficient (a measure of the degree of relatedness between the parents of a focal individual) or by comparing fitness estimates between experimentally inbred and outbred individuals. However, obtaining estimates of compound fitness is difficult, and consequently most studies use only some components of fitness or traits correlated with it. This can lead to severe underestimation of the magnitude of inbreeding depression (e.g., Hedrick and Garcia-Dorado 2016; Huisman et al. 2016; Stoffel et al. 2020). Moreover, in experimental studies, only individuals that survive until the time when fitness is assessed are included, which may be problematic in cases where inbreeding affects pre-measurement (e.g., juvenile) survival, and which again may lead to an underestimation of inbreeding depression (Ritland 1990; Willis 1993; Lande et al. 1994; Harrisson et al. 2019).

In the present study, we investigated how the magnitude of inbreeding depression accumulates across the entire life cycle of a cyclical parthenogenetic species. We minimized the possibility of selective pre-measurement mortality by taking an early embryonic stage as the starting point for our study. Furthermore, using a cyclical parthenogenetic species, in which several asexual generations follow a sexual one, we were able to monitor fitness not only of the sexually produced inbred and outbred individuals, but also of their asexual offspring. Indeed, while one generation of sexual outcrossing erases effects of previous inbreeding, clonal forms of asexuality maintain inbreeding levels (and other forms of asexuality, such as automixis, may even increase inbreeding levels, e.g., Svendsen et al. 2015). Thus, because inbred genotypes remain inbred across asexual generations, inbreeding depression may be an important factor in the evolution of partial asexual life cycles, a possibility that has so far received very little attention (Muirhead and Lande 1997; Marriage and Kelly 2009; Navascués et al. 2010; Vallejo-Marín et al. 2010).

We assessed inbreeding depression in the planktonic crustacean *Daphnia magna*, which inhabits diverse small to medium-sized water bodies across the northern hemisphere. Most populations of the species are intermittent: each year, only sexually-produced diapause stages survive harsh periods (summer drought or winter freezing). These diapause stages then found active populations when conditions for planktonic stages become favorable. The planktonic stages reproduce parthenogenetically throughout the favorable season, and bouts of parthenogenetic production of males (sex being evnironmentally determined) and sexual reproduction occur, mostly towards the end of the season. Sexual reproduction leads to the formation of new diapause stages, which will typically remain dormant until the next favorable season. While parthenogenetic reproduction is meiosis-derived (Hiruta et al. 2010), the genotype of the mother is transmitted with only very little changes (Dukić et al. 2019), so that, for the purpose of this study, we assumed clonal reproduction (i.e., no change in genotype nor inbreeding level during parthenogenetic reproduction).

We used an experimental approach combined with genetic markers. We cultured parental clones (parthenogenetic descendants of individual females obtained from the field) first in the laboratory and then in large cultures outdoors. Clones were paired randomly, while assuring that the two clones of each pair were homozygous for different alleles at one or two microsatellite loci (“diagnostic loci”). The clones of each pair were then placed together in the same mass culture, and sexually produced diapause stages were collected. These were either formed by inbreeding of either or the two parental clones (within-clone mating is genetically equivalent to self-fertilization) or by outcrossing. A sample of diapause stages was genotyped at the diagnostic loci to establish initial frequencies of inbred and outbred genotypes. The rest of the diapause stages were overwintered and exposed to hatching conditions, again in large outdoor containers, in the next spring. The hatchlings grew to adulthood and started reproducing, thus forming experimental populations consisting of inbred and outbred genotypes. These populations were grown under ambient conditions until the next autumn, and additional samples were taken at five time points to establish the frequencies of inbred and outbred genotypes (note that each sexually produced hatchling represents a genetically unique individuals, while clonal copies of these initial hatchlings exist later in the season, just as in natural *D. magna* populations).

## MATERIALS AND METHODS

### Origin of clones and establishment of clonal cultures

*Daphnia magna* clones used in the experiment were established from individual females isolated from a sample obtained in spring 2017 from a 0.13 ha pond in Moscow Zoo [N55.7635°, E37.5813°], the “source population”, shortly after establishment of the active population from diapause stages. Clones were cultured in the laboratory, in jars filled with 150 ml of *Daphnia* medium (Klüttgen et al. 1994). The cultures were kept at 18°C, in a 16:8 h light:dark photoperiod and fed three times a week with 2*10^5^ *Scenedesmus acutus* cells per ml of culture medium. Clones were genotyped at two microsatellite loci known to be polymorphic in this population (Reisser et al. 2017). Based on their genotypes, we selected four pairs of parent clones so that the clones within each pair were homozygous for different alleles at one or both loci (Table S1).

The eight selected parent clones were first transferred to 5-L aquaria, kept under the same laboratory conditions as detailed above. In late July, 2017, after they had grown in numbers, they were transferred to outdoor tanks, placed under ambient conditions in the garden of the Koltsov Institute of Developmental Biology, Moscow, at a few kilometers distance from the source population. Ten days before the outdoor cultures were initiated, each tank had been filled with 120 L tap water, 300 g of marble chips as calcium source, and 150 g of horse manure contained in bags made of 200 μm mesh net as fertilizer. Each tank was also inoculated with 10 liters of pond water from the source pond, filtered twice through a 200 μm mesh net to avoid contamination but still allow a natural inoculum of microalgae and bacteria. The tanks were then covered with mosquito nets and kept that way without addition of further food for the whole duration of the experiment. After the transfer, the clonal cultures were allowed to propagate for two months. On 25 September 2017, the tanks contained between 10’000 and 50’000 individuals each and showed a high degree of sexual reproduction (male and diapause egg production).

### Crosses

In order to obtain inbred and outbred sexual offspring, the cultures of the two parent clones per pair were mixed in late September 2017 and grown together in mass culture under ambient conditions for a further month (previously produced diapause stages were discarded at the time of mixing). For the mixed cultures, additional tanks were prepared as above (using only 90 L of water) and about 10000 individuals of each of the two parent clones were transferred to these new tanks. Two replicate mixed cultures were established for each of the four pairs (i.e., eight tanks in total). In the end of October 2017, 600-1000 ephippia containing dipausing embryos were collected from the bottom of each tank and transferred to 50-ml tubes, which were placed in a dark container, kept at 4°C for approximately 4.5 months of overwintering.

### Offspring populations and sampling

In mid-April 2018, a part of the diapause stages were opened, and, for each of the eight replicate crosses, 70-100 diapausing embryos were preserved in ethanol for later genotyping. Note that each diapause stage (“ephippium”) usually contains two such embryos, with arrested development at roughly a 3500 cell stage (Chen et al. 2018). The remaining diapause stages were removed from cold and dark diapause conditions and transferred to beakers with 150 ml of *Daphnia* medium, placed in a climatic chamber at 10°C and constant light. These conditions are known to stimulate diapause breaking in *D. magna* from the source population (Galimov et al. 2011). Hatching of juveniles started after approximately seven days. The beakers were checked regularly for new hatchlings. Every tenth hatchling was removed and preserved in ethanol for later genotyping. The remaining hatchlings were transferred to jars with 50 ml of *Daphnia* medium per hatchling, placed in the same climatic chamber and fed every day with 4×10^4^ *S. acutus* cells per ml of medium. Five days after hatchlings, the juveniles from each replicate were transferred to a new outdoor tank, which had been prepared as described above for the 2017 mass cultures of the parental clones. In total, we thus established eight outdoor experimental populations, each consisting of a mixture of inbred and outbred genotypes from one pair of parental clones (two replicate populations per pair). Each population was founded with 1020 to 2500 juveniles, depending on the number of hatchlings obtained for each replicate. Three of the populations were started on 20 April 2018 and five populations on April 28 2018. Each population was sampled for genotyping on four further dates: 17 May, 4 June, 18 July, and 7 September. Depending on the population size, 1/20 or 1/10 of the water volume was taken from each tank after thorough mixing and filtered through a 200 μm mesh net to remove all *Daphnia* from the sampled volume. All removed *Daphnia* were preserved in ethanol, kept at −20°C until genotyping. An additional sample on 7 May, mainly consisting of first-generation parthenogenetic offspring of the hatchlings, was obtained from three replicate cultures (03×56A, 08×17B, and 18×90A) and used for some analyses (sex ratios, brood sizes, genotypes of males). For adult females and prior to genotyping, we distinguished females carrying parthenogenetic broods, females carrying ephippial broods, and females with an empty brood pouch, by inspecting samples under a stereomicroscope. For females carrying a parthenogenetic brood, we counted the number of eggs to determine the brood size. Finally, we determined adult sex ratios by counting the number of adult males and adult females in a larger sample than the one that was genotyped (eleven samples from six replicate populations, Table S2),

### Genotyping

Total genomic DNA was extracted from whole *Daphnia* or whole embryos using Glass Fiber Plate DNA Extraction (Ivanova et al. 2006). Except for embryos and hatchlings, genotyping was mainly carried out on adult females. Generally, we aimed for sample sizes of at least 64 individuals per sample (see Table S3 for all sample sizes). Where sample sizes of adult females were insufficient, pre-adult females (i.e., individuals being 1-3 days from reaching adulthood) were added. For females carrying ephippial broods, we removed the sexually produced embryos contained in these ephippia prior to DNA extraction in order to avoid contamination with embryonal DNA. The genotypes of males were analyzed in five samples (Table S4).

### Microsatellite analysis

We used the primers described in Reisser et al. (2017), with forward primers being labelled with R6G (locus STR50) or FAM (locus STR050). For polymer chain reactions we used Evrogen PCR kits with Hot Start thermostable polymerase (HS Taq polymerase, Evrogen, Russia). PCR reactions were carried out for each locus separately in a total volume of 14.1 μl (0.2 μl of a primer mixture containing forward and reverse primers, each at 10mkM, 0.15 μl of HS Taq polymerase, 1.5 μl of 10Х Taq Turbo buffer, 0.25 μl dNTP, and 12 μl bidistilled water). PCR started with an initial denaturation at 95°C (4 min), followed by four cycles of 95°C (40 sec), 50°C (1 min 30 sec), 70°C (1 min) and then by 35 cycles with annealing temperature of 52°C instead of 50°C. This was followed by a final extension (5 min at 70°C), after which the reactions were put on hold at 4°C. The PCR products of the two loci were then mixed in equal proportions (1.5 μl each) and diluted with 48.5 μl distilled water. Of this mixture, 1.5 μl were used for electrophoresis, the remaining part was stored at −20°C as backup. A formamide solution (Hi-Di) was used as a size standard during electrophoresis, and the results of the electrophoresis were analyzed with the software GeneMarker (SoftGenetics, USA).

### Statistical analysis

For the first sample (embryos), we tested for each of the replicates whether genotype frequencies were in Hardy-Weinberg equilibrium, using a randomization test implemented in FSTAT (Goudet 2003). To test whether starting conditions were similar between the two replicates of a pair, we tested whether the pair (here treated as a fixed effect) had a significant effect on the relative frequencies of inbred of one vs. the other parent, and on the relative frequencies of inbred genotypes (from both parents combined) vs. outbred genotypes. This was done using generalized linear models, assuming binomial error distribution (each individual belonged to either of two categories, the more common vs. the rarer class of inbred genotypes in the first model, inbred vs. outbred genotypes in the second model). The models were evaluated using a maximum likelihood approach with Firth’s bias correction and with a logit link function. We used *χ*^*2*^-tests to evaluate if frequencies of inbred genotypes (inbred offspring from both parents combined) vs. outbred genotypes changed within replicate cultures (and Fisher’s exact tests for comparisons that were made only between two samples). To analyze the frequency changes of inbred vs. outbred genotypes across replicate cultures, we first calculated the difference in the estimated frequencies within each replicate for the two samples in question, then calculated the average of this difference for each of the four pairs, and then tested whether the arithmetic mean of those four averages was significantly different from zero, using a one-sample *t*-test. This may not be the most powerful test for the comparison of subsequent samples, but it accurately reflects the fact that the two replicates of a pair were non-independent, is simple, and only makes a limited number of assumptions that are likely to be approximately met (frequency difference may not very well follow a normal distribution, but the means of pair means should meet the normal distribution assumption sufficiently well for *P*-values evaluated with one-sample *t*-test to be highly accurate).

We also estimated the magnitude of inbreeding depression *δ* = 1 - *w*_*in*_/*w*_*out*_, where *w*_*in*_/*w*_*out*_ is the relative fitness of inbred to outbred genotypes, estimated from the frequency changes according to ln(*w*_*in*_/*w*_*out*_) = ln(*in*_*t*_/*out*_*t*_) - ln(*in*_*0*_/*out*_*0*_), where *in*_*t*_ and *out*_*t*_ and *in*_*0*_ *out*_*0*_ are the frequencies of inbred genotypes (from both parents combined) and outbred genotypes at times *t* and zero, respectively. This provides an estimate of inbreeding depression per unit of time (Johnston and Schoen 1994; Haag et al. 2002) which, in our case, was taken to be either the entire experiment (145 days). Similarly, we estimated the diploid load of lethal equivalents 2*B* according to 2*B* = –2(ln(*w*_*in*_/*w*_*out*_))/*F*, with *F* = 0.5 being the inbreeding coefficient (Hedrick and Garcia-Dorado 2016; Nietlisbach et al. 2019). The mean estimates of *δ* and 2*B* were computed from the estimates of the average frequency of inbreds (inbred offspring from both parents combined) in samples 1 and 6 across all pairs (means of pair means). To obtain confidence limits, we estimated *δ* and 2*B* for each replicate culture separately. For the four replicate cultures, in only outbred genotypes were observed in sample 6, we assumed that a sample of twice the actual sample size would have contained one inbred genotype. We then estimated the 95 % confidence interval from the standard error of pair means of these estimates.

## RESULTS

The eight experimental populations exhibited seasonal cycles that were similar to the typical seasonal cycle in the natural source population (Galimov et al. 2011), with a short period of fast parthenogenetic reproduction in the beginning of the season and a subsequent strong decline in fecundity and population growth (Fig. S1, Table S2). In the natural source population, the production of ephippia (i.e., diapause stages containing sexually-produced embryos) and males is almost entirely limited to the end of the season, except for a small peak of male production and (very rarely) ephippia production occurs in late spring (Galimov et al. 2011). This small early peak occurred also in our experiment, but was somewhat more pronounced than in the natural source population (Fig. S1, Table S2). Nonetheless, just as in the natural population, the vast majority of sexual reproduction in the parent generation occurred in autumn and would therefore have occurred in the offspring generation after the experiment was terminated.

In the first sample (embryos), outbred genotypes as well as both types of inbred genotypes (i.e., inbred offspring of both parental clones) were present in all eight replicate cultures. Genotype frequencies were similar between the two replicates of a pair. The “pair” effect on the relative frequencies of the more common *vs*. the rarer class of inbred genotypes was not significant (*χ*^*2*^ = 7.1, *d.f.* = 3, *P* = 0.069) but within each pair, it was always the inbred offspring of the same parent that were more abundant in both replicates. This likely reflects fitness differences between the two parental clones in the previous autumn (having led to similar frequency differences in both replicates of a given pair) or differences in investment into sexual reproduction. Moreover, there was a significant “pair” effect on the relative frequencies of outbred *vs*. all inbred genotypes from both parents combined, *χ*^*2*^ = 66.8, *d.f.* = 3, *P* < 0.0001). In six out of the eight replicate cultures, there was a slight to rather strong excess of inbred genotypes compared to Hardy-Weinberg expectations (positive *F*_*IS*_, significant in three replicate cultures, Table S3). The remaining two replicates showed a slight to moderate excess of outbred genotypes (negative *F*_*IS*_, significant in one replicate culture, Table S3).

Already in the second sample (hatchlings), the frequency of inbred genotypes (from both parents combined) showed a decrease in six of the eight replicate cultures (significant Fisher’s exact test in three cultures, Fig. 1, Table S3). Overall, across replicate cultures, and considering the pair as the unit of replication, the frequency of inbred genotypes was 12.7 % lower among hatchlings than among embryos (*t* = −7.8, df = 3, *P* = 0.004). The frequency of inbred genotypes then showed a more or less steady further decrease over subsequent samples. Only between mid-May and early June (between sample 3 and sample 4), there was a slight, but non-significant increase in the frequency of inbred genotypes (*t* = 0.31, df = 3, *P* = 0.78). All other pairwise comparisons between subsequent samples were significant (*P* < 0.05), except between sample 5 and sample 6 (*t* = −1.39, df = 3, *P* = 0.26) At the end of the experiment, almost no inbred genotypes were left in any of the replicate cultures (Fig. 1). The decrease in the frequency of inbred genotypes from the first to the last sample was highly significant (*t* = −8.7, df = 3, *P* = 0.003; within-replicate population Fisher’s exact tests, *P* < 0.0001 in all eight replicate populations). The mean estimate of inbreeding depression *δ* for the entire experiment was 0.992 (95 % confidence limits: 0.979, 1), which translates to a diploid load of lethal equivalents 2*B* for survival of a genotype to the end of the experiment of 19.4 (95 % confidence limits: 16.5, 22.9).

**Fig 1.**
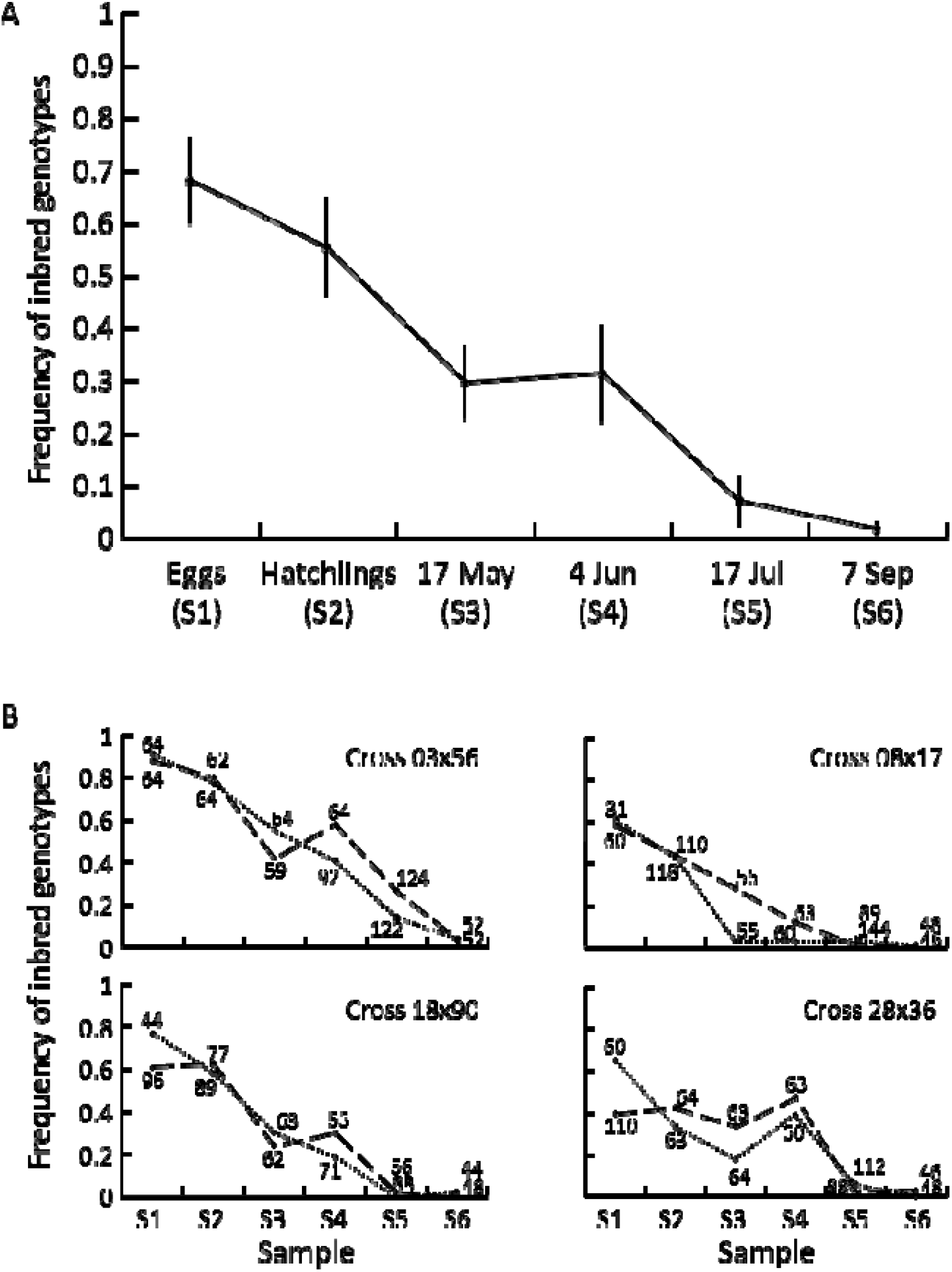
A) Average frequency of inbred genotypes at each sampling date and across all replicate populations (error bars indicate standard errors). B) Frequencies of inbred genotypes in each replicate population. The four panels depict the four parental crosses (replicate A: dotted line, replicate B: hatched line) and numbers next to each data point represent the number of genotyped individuals.

Genotypes of males were obtained for only five samples (Table S4). In one of them, there was only a single male, which was outbred and in another one, the frequency of inbred genotypes was slightly, but non-significantly higher among males than among females (Fisher’s exact test, *P* = 0.30, Table S4). However, in the remaining three samples, the frequency of inbred genotypes was significantly lower in males than in females (Fisher’s exact tests, *P* < 0.006, Table S4), indicating that inbred genotypes invested less in male production than outbred genotypes.

Regarding adult females (Table S5), there were no significant differences in parthenogenetic brood sizes between inbred and outbred females (paired *t* = −1.11, d.f. = 15, *P* = 0.29) nor in the proportion of inbred genotypes among females with empty brood pouches (zero broods, paired *t* = −1.41, d.f. = 24, *P* = 0.17) nor among ephippial females (paired *t* = 1.79, d.f. = 7, *P* = 0.12), when compared to the frequency of inbred genotypes among all other females. However, if anything, there was a tendency for inbred females to produce less ephippia than outbred females (Fisher’s exact test, *P* = 0.09 in one of the eight samples in which this could be analyzed, and paired *t* = 2.18, d.f. = 7, *P* = 0.07 across the eight samples, with samples weighted according to the overall frequency of inbred genotypes to account for the low statistical power in samples where inbred genotypes were rare). Inbred females also showed a tendency to more often have empty brood pouches compared to outbred females (Fisher’s exact test, *P* < 0.05 in five of the 25 samples in which this could be evaluated, Table S5).

## DISCUSSION

We found very strong inbreeding depression in *D. magna* when following offspring of within-clone matings (genetically equivalent to self-fertilization) and outcrossed offspring in common outdoor cultures for an entire season. Previous estimates of the magnitude of inbreeding depression in *D. magna*, and other *Daphnia* species were mostly based on individual life-history traits but also on the intrinsic rate of population growth and frequency changes during competition (De Meester 1993; Deng 1997; Deng and Lynch 1997; Salathé and Ebert 2003; Lohr and Haag 2015). These estimates were generally also high but the estimates of the present study largely exceed all previous estimates, including for clones from the same source population as used here (Lohr and Haag 2015). The estimate of the diploid number of equivalents is very high, also in comparison with a large range of other organisms (Hedrick and Garcia-Dorado 2016; Nietlisbach et al. 2019). The strong inbreeding depression across the entire experiment occurred due to gradual and cumulative decreases in the frequency of inbred genotypes, already starting during hatching and then maintained during the whole season with a possible exception between mid-May and early June (which is possibly explained by slower growth of inbred individuals leading to an underestimation of their frequency in mid-May, as only adults were genotyped). It is likely that inbreeding affected several fitness-related traits (e.g., hatching, fecundity, and survival), and the gradual decrease in the frequencies of inbred genotypes reflected the sum of these components. Overall, our results are thus in line with findings on other organisms suggesting that the magnitude of inbreeding depression can be strongly underestimated if estimates of inbreeding depression based on components or correlates of fitness only (e.g., Hedrick and Garcia-Dorado 2016; Huisman et al. 2016; Stoffel et al. 2020). Moreover, the observation of reduced frequency of inbred genotypes already among hatchlings and even more so in sample 3 suggests that substantial inbreeding depression may occur already before the time point when a typical life table experiment would be carried out (e.g., Lohr and Haag 2015). Such pre-experimental inbreeding depression may further contribute to underestimation of inbreeding depression in classical experiments (Ritland 1990; Willis 1993; Lande et al. 1994; Harrisson et al. 2019).

In the natural source population, sexual reproduction takes place almost exclusively in the end of the season (Galimov et al. 2011), and hence the population of the following year will almost entirely be produced by clones surviving to until then. Assuming that the results from our outdoor cultures can be extrapolated to the natural population, we conclude that the fitness of genotypes produced by within-clone mating is very close to zero because they have a very low likelihood to survive until sexual reproduction and thus to contribute to the population of the following year. Note also that within-clone mating is the form of inbreeding most likely to occur in *Daphnia* because clonal selection can lead to situations where some clones (i.e., the parthenogenetic descendants of a single hatchling at the start of the season) reach high frequencies in the end of the season, so that within-clone mating may occur even under random mating (Pfrender and Lynch 2000; De Meester et al. 2006; Vanoverbeke and De Meester 2010; Hamrová et al. 2011). The outdoor cultures did differ from the natural population in terms of their earlier investment into sexual reproduction (Galimov et al. 2011). However, our results do not suggest that early investment into sexual reproduction has contributed to the decrease in the frequency of inbred genotypes: While investment into males and ephippia is known to slow clonal growth in *Daphnia* because they trade off with parthenogenetic production of female offspring (e.g., Gerber et al. 2018), male and ephippia production did not differ between inbred and outbred genotypes (or there was an inverse tendency, potentially indicating additional inbreeding depression in these traits). Finally, not all *Daphnia* populations are as strictly seasonal as the source population and inbreeding depression is also known to vary substantially among populations, with the source population being on the higher end (Lohr and Haag 2015). Nonetheless, taken together, our results suggest that the production of offspring by within-clone mating entails high costs in *Daphnia* and that these costs are accentuated by asexual reproduction.

On a more general level, our study suggests that inbreeding depression may be an underappreciated issue in organisms that can undergo both sexual and asexual reproduction (i.e., in partial asexuals). Besides the fact that inbred genotypes remain inbred across several asexual generations, asexuality may also increase the likelihood of matings to occur between highly similar genotypes compared to fully sexual organisms (e.g., Haag et al. 2005; Thompson et al. 2008). However, it is unclear, how effects of partial asexuality would play out on the long run, as both the increased likelihood of producing inbred offspring and the reduced effective population sizes of partially asexual organisms (compared to fully sexual ones) may lead to purging of deleterious mutations (Glémin 2003; Marriage and Kelly 2009; Hartfield 2016). The few theoretical and empirical studies on inbreeding depression in partial asexuals mostly concentrated on plant life histories with intermittent vegetative growth and on effects of somatic mutations (Muirhead and Lande 1997; Marriage and Kelly 2009; Navascués et al. 2010; Vallejo-Marín et al. 2010; Marriage and Orive 2012). Our results demonstrate that the negative fitness consequences of inbreeding can accumulate across asexual generations and result in very high overall levels of inbreeding depression (i.e., almost zero contribution of inbred genotypes to future sexual generations). While further studies are needed to determine how partial asexuality generally interacts with inbreeding depression, our findings intuitively suggest that evolving a partially asexual life cycle from a purely sexual one may entail costs in terms of elevated inbreeding depression, at least initially. They also suggest that partial asexuality may favor the evolution of self-incompatibility (Navascués et al. 2010; Vallejo-Marín et al. 2010) or other inbreeding-avoidance mechanisms. Interestingly, some genotypes of *D. magna* (not included in the present study) have indeed evolved obligate outcrossing: They do not produce males and participate in sexual reproduction only as females (so-called non-male producing genotypes, Galimov et al. 2011; Reisser et al. 2017; Molinier et al. 2019). To our knowledge, these genotypes only occur in populations with strong inbreeding depression, and their evolution and equilibrium frequency is indeed predicted to be influenced by both the magnitude of inbreeding depression and the frequency of within-clone mating in genotypes that do produce males (Innes and Dunbrack 1993). Clearly, inbreeding depression may have a strong impact on evolutionary processes in partial asexuals and this may warrant further study.

## Acknowledgments

We thank Camille Ameline, Maridel Fredericksen, Anastasia Raeva and Aleksey Popov for their help with sampling and experiments. VGT, AAZ and YRG were supported by Grant 16-04-01579 from Russian Foundation for Basic Research to YRG and by the Assignment of Ministry of Education and Science of the Russian Federation to IDB RAS (Institute for Developmental Biology of the Russian Academy of Sciences). CRH was supported by Grant ANR-17-CE02-0016-01, GENASEX, from the Agence Nationale de la Recherche. The project used the equipment of IDB RAS Core Centrum.

## SUPPLEMENTARY MATERIALS

**Fig. S1.**
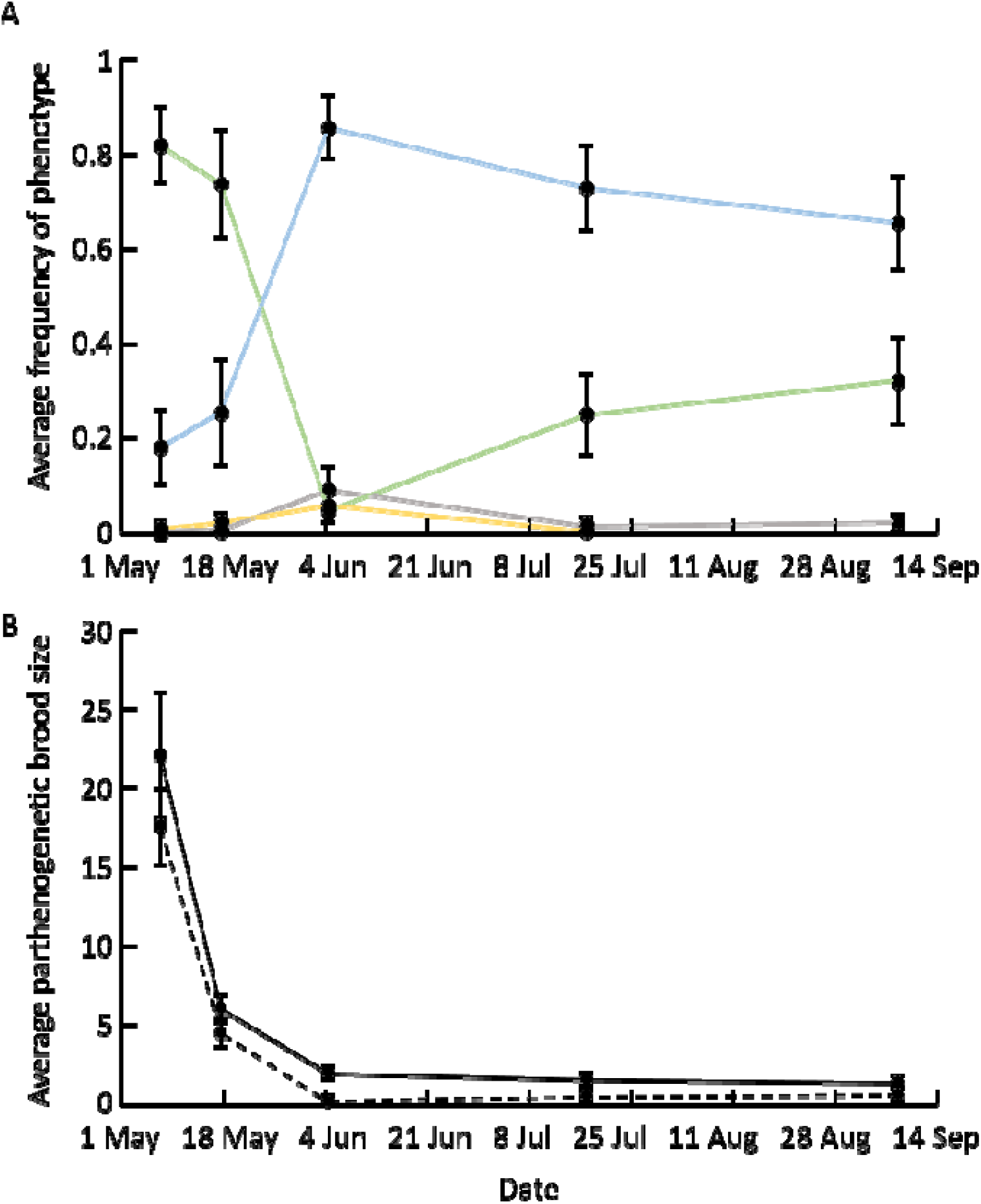
A) Seasonal changes in the average frequency of females with parthenogenetic broods (green line), females with empty brood pouches (blue line), and females with ephippial broods (grey line) among all adult females. In addition, the average male-to-female sex ratio is shown (yellow line, four dates only), the estimation of which was based on larger population samples (see Table S2). B) Seasonal changes in average parthenogenetic brood sizes of non-zero parthenogenetic broods (solid line) and when including brood sizes of zero (i.e., including females with empty brood pouches, hatched line). Error bars indicate standard errors across replicate cultures.

**Table S1.** Pairs of parental clones and their diagnostic microsatellite genotypes. Locus names as in Reisser et al. (2017).

| Cross | First parent clone |  |  | Second parent clone |  |  |
| --- | --- | --- | --- | --- | --- | --- |
|  | Clone name | Genotype |  | Clone name | Genotype |  |
|  |  | Locus STR50 | Locus STR74 |  | Locus STR50 | Locus STR74 |
| 03x56 | Clone_03 | 238/238 | 190/190 | Clone_56 | 232/232 | 192/192 |
| 08x17 | Clone_08 | 232/232 | 190/190 | Clone_17 | 238/238 | 190/192 |
| 18x90 | Clone_18 | 238/238 | 190/190 | Clone_90 | 232/232 | 192/192 |
| 28x36 | Clone_28 | 232/232 | 190/190 | Clone_36 | 244/244 | 190/190 |

**Table S2.** Seasonal dynamics of proportion of adult females with different brood types (parthenogenetic broods, empty brood pouches, ephippial broods), average brood sizes (estimated for all non-zero parthenogenetic broods and for all non-ephippial females), and adult male-to-female sex ratios. Female phenotypes and brood sizes were based on a different sample (*N* ad. ♀) than the adult sex ration (*N* for ad. sex ratio).

| Cross and replicate culture | Date | $N$ ad. ♀ | Proportion of adult females with | | | Avg. non-zero brood size | Avg. brood size incl. zeros | $N$ for ad. sex ratio | Adult sex ratio |
| --- | --- | --- | --- | --- | --- | --- | --- | --- | --- |
|  |  |  | Parth. broods | No broods | Ephippial broods |  |  |  |  |
| 03x56A | 7 May | 64 | 0.95 | 0.05 | 0 | 14.37 | 13.7 |  |  |
| 03x56A | 17 May | 64 | 0 | 1 | 0 | NA | 0 |  |  |
| 03x56A | 4 Jun | 64 | 0 | 1 | 0 | NA | 0 | 1068 | 0.1 |
| 03x56A | 17 Jul | 122 | 0.29 | 0.66 | 0.05 | 1.63 | 0.49 |  |  |
| 03x56A | 7 Sep | 52 | 0.04 | 0.96 | 0 | 1 | 0.04 |  |  |
| 03x56B | 17 May | 59 | 0.88 | 0.12 | 0 | 6.38 | 5.62 |  |  |
| 03x56B | 4 Jun | 64 | 0.11 | 0.61 | 0.28 | 2.57 | 0.39 |  |  |
| 03x56B | 17 Jul | 124 | 0.13 | 0.87 | 0 | 1.25 | 0.16 |  |  |
| 03x56B | 7 Sep | 24 | 0.04 | 0.92 | 0.04 | 1 | 0.04 |  |  |
| 08x17A | 17 May | 55 | 0.91 | 0.09 | 0 | 4.80 | 4.36 | 674 | 0.01 |
| 08x17A | 4 Jun | 60 | 0 | 0.93 | 0.07 | NA | 0 | 1204 | 0.07 |
| 08x17A | 17 Jul | 89 | 0 | 1 | 0 | NA | 0 |  |  |
| 08x17A | 7 Sep | 48 | 0.54 | 0.46 | 0 | 1.58 | 0.86 |  |  |
| 08x17B | 7 May | 45 | 0.69 | 0.31 | 0 | 25.30 | 17.42 | 213 | 0 |
| 08x17B | 17 May | 56 | 0.82 | 0.18 | 0 | 3.87 | 3.18 | 508 | 0 |
| 08x17B | 4 Jun | 63 | 0.05 | 0.95 | 0 | 1.67 | 0.08 | 393 | 0.01 |
| 08x17B | 17 Jul | 58 | 0.02 | 0.98 | 0 | 1 | 0.02 |  |  |
| 08x17B | 7 Sep | 46 | 0.30 | 0.70 | 0 | 1.29 | 0.39 |  |  |
| 18x90A | 7 May | 43 | 0.81 | 0.19 | 0 | 27.12 | 22.07 | 310 | 0.02 |
| 18x90A | 17 May | 63 | 0.73 | 0.22 | 0.05 | 3.35 | 2.57 | 1215 | 0.07 |
| 18x90A | 4 Jun | 71 | 0 | 1 | 0 | NA | 0 |  |  |
| 18x90A | 17 Jul | 40 | 0.4 | 0.5 | 0.10 | 2.389 | 1.06 |  |  |
| 18x90A | 7 Sep | 44 | 0.66 | 0.20 | 0.14 | 2.321 | 1.77 |  |  |
| 18x90B | 17 May | 62 | 0.97 | 0.03 | 0 | 7.21 | 6.98 |  |  |
| 18x90B | 4 Jun | 53 | 0 | 0.91 | 0.09 | NA | 0 |  |  |
| 18x90B | 17 Jul | 13 | 0.46 | 0.54 | 0 | 1.33 | 0.61 |  |  |
| 18x90B | 7 Sep | 48 | 0.52 | 0.48 | 0 | 1.44 | 0.75 |  |  |
| 28x36A | 17 May | 64 | 0.83 | 0.17 | 0 | 9.11 | 7.55 | 386 | 0.02 |
| 28x36A | 4 Jun | 60 | 0.17 | 0.53 | 0.30 | 2.20 | 0.52 |  |  |
| 28x36A | 17 Jul | 31 | 0.65 | 0.35 | 0 | 2.50 | 1.61 |  |  |
| 28x36A | 7 Sep | 48 | 0.06 | 0.94 | 0 | 1 | 0.06 |  |  |
| 28x36B | 17 May | 63 | 0.76 | 0.24 | 0 | 7.92 | 6.03 |  |  |
| 28x36B | 4 Jun | 63 | 0.08 | 0.92 | 0 | 1.40 | 0.11 | 580 | 0.05 |
| 28x36B | 17 Jul | 88 | 0.08 | 0.92 | 0 | 1 | 0.08 | 171 | 0.006 |
| 28x36B | 7 Sep | 46 | 0.41 | 0.59 | 0 | 1.47 | 0.61 |  |  |

**Table S3.** Number of inbred offspring of each parent of a given cross (inbred1, inbred2), number of outbred offspring, and *F*_*IS*_ in each sample of each replicate population (A, B) of each cross. The crosses are defined by the clone names of the parent clones according to Table S1. Sampling time points (“Sample”) are defined by sampling date/stage according to Fig. 1 in the main text.

| Sampling date/stage | Sample | Cross and replicate culture | $N$ inbred1 | $N$ inbred2 | $N$ outbred | $F_{IS}$ |
| --- | --- | --- | --- | --- | --- | --- |
| Eggs | S1 | 03x56A | 46 | 13 | 5 | 0.79 |
| Hatchlings | S2 | 03x56A | 49 | 1 | 14 | 0 |
| 17 May 2018 | S3 | 03x56A | 36 | 0 | 28 | -0.28 |
| 04 Jun 2018 | S4 | 03x56A | 38 | 0 | 54 | -0.42 |
| 17 Jul 2018 | S5 | 03x56A | 15 | 3 | 104 | -0.72 |
| 07 Sep 2018 | S6 | 03x56A | 1 | 1 | 50 | -0.92 |
| Eggs | S1 | 03x56B | 54 | 3 | 7 | 0.41 |
| Hatchlings | S2 | 03x56B | 49 | 1 | 12 | 0.03 |
| 17 May 2018 | S3 | 03x56B | 21 | 4 | 34 | -0.26 |
| 04 Jun 2018 | S4 | 03x56B | 31 | 6 | 27 | 0 |
| 17 Jul 2018 | S5 | 03x56B | 33 | 1 | 90 | -0.56 |
| 07 Sep 2018 | S6 | 03x56B | 1 | 1 | 50 | -0.92 |
| Eggs | S1 | 08x17A | 17 | 2 | 12 | 0.006 |
| Hatchlings | S2 | 08x17A | 45 | 6 | 67 | -0.27 |
| 17 May 2018 | S3 | 08x17A | 2 | 0 | 53 | -0.93 |
| 04 Jun 2018 | S4 | 08x17A | 2 | 0 | 58 | -0.94 |
| 17 Jul 2018 | S5 | 08x17A | 2 | 2 | 85 | -0.91 |
| 07 Sep 2018 | S6 | 08x17A | 0 | 0 | 48 | -1 |
| Eggs | S1 | 08x17B | 34 | 1 | 25 | -0.19 |
| Hatchlings | S2 | 08x17B | 38 | 10 | 62 | -0.21 |
| 17 May 2018 | S3 | 08x17B | 12 | 4 | 40 | -0.46 |
| 04 Jun 2018 | S4 | 08x17B | 3 | 5 | 55 | -0.75 |
| 17 Jul 2018 | S5 | 08x17B | 2 | 0 | 142 | -0.97 |
| 07 Sep 2018 | S6 | 08x17B | 0 | 0 | 46 | -1 |
| Eggs | S1 | 18x90A | 28 | 6 | 10 | 0.4 |
| Hatchlings | S2 | 18x90A | 48 | 4 | 37 | -0.1 |
| 17 May 2018 | S3 | 18x90A | 17 | 2 | 44 | -0.48 |
| 04 Jun 2018 | S4 | 18x90A | 12 | 1 | 58 | -0.67 |
| 17 Jul 2018 | S5 | 18x90A | 0 | 0 | 88 | -1 |
| 07 Sep 2018 | S6 | 18x90A | 1 | 0 | 43 | -0.96 |
| Eggs | S1 | 18x90B | 49 | 10 | 37 | 0.08 |
| Hatchlings | S2 | 18x90B | 43 | 5 | 29 | 0 |
| 17 May 2018 | S3 | 18x90B | 15 | 0 | 47 | -0.61 |
| 04 Jun 2018 | S4 | 18x90B | 14 | 2 | 37 | -0.47 |
| 17 Jul 2018 | S5 | 18x90B | 1 | 0 | 55 | -0.96 |
| 07 Sep 2018 | S6 | 18x90B | 0 | 0 | 48 | -1 |
| Eggs | S1 | 28x36A | 6 | 33 | 21 | 0.13 |
| Hatchlings | S2 | 28x36A | 3 | 18 | 42 | -0.41 |
| 17 May 2018 | S3 | 28x36A | 0 | 11 | 53 | -0.71 |
| 04 Jun 2018 | S4 | 28x36A | 3 | 20 | 37 | -0.34 |
| 17 Jul 2018 | S5 | 28x36A | 0 | 6 | 106 | -0.9 |
| 07 Sep 2018 | S6 | 28x36A | 0 | 0 | 48 | -1 |
| Eggs | S1 | 28x36B | 11 | 32 | 67 | -0.26 |
| Hatchlings | S2 | 28x36B | 5 | 22 | 37 | -0.24 |
| 17 May 2018 | S3 | 28x36B | 1 | 20 | 42 | -0.47 |
| 04 Jun 2018 | S4 | 28x36B | 8 | 22 | 33 | -0.1 |
| 17 Jul 2018 | S5 | 28x36B | 0 | 1 | 87 | -0.98 |
| 07 Sep 2018 | S6 | 28x36B | 0 | 1 | 45 | -0.96 |

**Table S4.** Proportion of inbred genotypes in males vs. females.

| Cross and replicate culture | Sampling date | Females |  | Males |  | *P* Fisher's exact |
| --- | --- | --- | --- | --- | --- | --- |
|  |  | *N* | Proportion inbred | *N* | Proportion inbred |  |
| 03x56A | 04 Jun 2018 | 92 | 0.41 | 55 | 0.51 | 0.30 |
| 08x17B | 17 Jul 2018 | 144 | 0.014 | 1 | 0 | 1 |
| 18x90A | 07 May 2018 | 63 | 0.43 | 64 | 0.094 | <0.0001 |
| 18x90A | 17 May 2018 | 63 | 0.3 | 59 | 0.068 | 0.001 |
| 18x90A | 04 Jun 2018 | 71 | 0.18 | 63 | 0.032 | 0.006 |

**Tables S5.** Proportion of inbred genotypes among females with empty brood pouches (“zero broods”), females with parthenogenetic broods, and females with ephippial broods, as well as average brood sizes of non-zero, parthenogenetic broods. Stars indicate a significant over-representation of inbred genotypes compared to all other females (* *P* < 0.05, ** *P* < 0.0). NA indicates that a proportion of average brood size was not assessed in a given sample. Note that the data in this table concern only adult females, which is the reason for the differences in numbers between this table and Table S3 (the latter includes sub-adult females in some cases).

| Cross and replicate culture | Sampling date | Females with zero broods |  | Females with parthenogenetic broods |  | Females with ephippial broods |  | Avg. (non-zero) brood size |  |
| --- | --- | --- | --- | --- | --- | --- | --- | --- | --- |
|  |  | N | Prop. inbred | N | Prop. inbred | N | Prop. inbred | Inbred | Outbred |
| 03x56A | 07 May 18 | 3 | 0.67 | 61 | 0.77 | 0 | NA | 15.06 | 12.07 |
| 03x56A | 17 May 18 | 64 | 0.56 | 0 | NA | 0 | NA | NA | NA |
| 03x56A | 04 Jun 18 | 64 | 0.34 | 0 | NA | 0 | NA | NA | NA |
| 03x56A | 17 Jul 18 | 81 | 0.19 | 35 | 0.09 | 6 | 0 | 1 | 1.69 |
| 03x56A | 07 Sep 18 | 50 | 0.04 | 2 | 0 | 0 | NA | NA | 1 |
| 03x56B | 17 May 18 | 7 | 0.57 | 52 | 0.4 | 0 | NA | 6.86 | 6.06 |
| 03x56B | 04 Jun 18 | 39 | 0.69* | 7 | 0.43 | 18 | 0.39 | 1.67 | 3.25 |
| 03x56B | 17 Jul 18 | 108 | 0.31** | 16 | 0 | 0 | NA | NA | 1.25 |
| 03x56B | 07 Sep 18 | 22 | 0.045 | 1 | 0 | 1 | 0 | NA | 1 |
| 08x17A | 17 May 18 | 5 | 0.2 | 50 | 0.02 | 0 | NA | 8 | 4.73 |
| 08x17A | 04 Jun 18 | 56 | 0.036 | 0 | NA | 4 | 0 | NA | NA |
| 08x17A | 17 Jul 18 | 89 | 0.045 | 0 | NA | 0 | NA | NA | NA |
| 08x17A | 07 Sep 18 | 22 | 0 | 26 | 0 | 0 | NA | NA | 1.58 |
| 08x17B | 07 May 18 | 14 | 0.79* | 31 | 0.42 | 0 | NA | 21.46 | 28.06 |
| 08x17B | 17 May 18 | 10 | 0.2 | 46 | 0.3 | 0 | NA | 4.57 | 3.56 |
| 08x17B | 04 Jun 18 | 60 | 0.12 | 3 | 0.33 | 0 | NA | 2 | 1.5 |
| 08x17B | 17 Jul 18 | 57 | 0.018 | 1 | 0 | 0 | NA | NA | 1 |
| 08x17B | 07 Sep 18 | 32 | 0 | 14 | 0 | 0 | NA | NA | 1.29 |
| 18x90A | 07 May 18 | 8 | 0.5 | 35 | 0.46 | 0 | NA | 22.38 | 31.11 |
| 18x90A | 17 May 18 | 14 | 0.21 | 46 | 0.35 | 3 | 0 | 3.06 | 3.5 |
| 18x90A | 04 Jun 18 | 71 | 0.18 | 0 | NA | 0 | NA | NA | NA |
| 18x90A | 17 Jul 18 | 20 | 0 | 16 | 0 | 4 | 0 | NA | 2.389 |
| 18x90A | 07 Sep 18 | 9 | 0.11 | 29 | 0 | 6 | 0 | NA | 2.321 |
| 18x90B | 17 May 18 | 2 | 0.5 | 60 | 0.23 | 0 | NA | 6.14 | 7.54 |
| 18x90B | 04 Jun 18 | 48 | 0.29 | 0 | NA | 5 | 0.4 | NA | NA |
| 18x90B | 17 Jul 18 | 7 | 0 | 6 | 0 | 0 | NA | NA | 1.33 |
| 18x90B | 07 Sep 18 | 23 | 0 | 25 | 0 | 0 | NA | NA | 1.44 |
| 28x36A | 17 May 18 | 11 | 0.091 | 53 | 0.19 | 0 | NA | 6.9 | 9.63 |
| 28x36A | 04 Jun 18 | 32 | 0.38 | 10 | 0.4 | 18 | 0.39 | 2 | 2.33 |
| 28x36A | 17 Jul 18 | 11 | 0 | 20 | 0.05 | 0 | NA | 4 | 2.42 |
| 28x36A | 07 Sep 18 | 45 | 0 | 3 | 0 | 0 | NA | NA | 1 |
| 28x36B | 17 May 18 | 15 | 0.67** | 48 | 0.23 | 0 | NA | 7.36 | 8.08 |
| 28x36B | 04 Jun 18 | 58 | 0.45 | 5 | 0.8 | 0 | NA | 1.25 | 2 |
| 28x36B | 17 Jul 18 | 81 | 0.012 | 7 | 0 | 0 | NA | NA | 1 |
| 28x36B | 07 Sep 18 | 27 | 0.037 | 19 | 0 | 0 | NA | NA | 1.47 |

